# Microtubule lattice defects facilitate spastin-mediated severing

**DOI:** 10.1101/2025.08.29.673059

**Authors:** Cordula Reuther, Paula Santos-Otte, Rahul Grover, Till Korten, Stefan Diez

## Abstract

The length regulation of microtubules and their organization into complex arrays inside cells occurs through the activity of polymerases, depolymerases as well as severing enzymes such as katanin and spastin. The latter hexamerize on the microtubule lattice, pull out single tubulin dimers in an ATP-dependent manner and eventually generate internal breaks in the microtubule. While both, katanin and spastin, were shown to be regulated by posttranslational tubulin modifications, only katanin was reported to have microtubule lattice-defect- or crossover-sensing activity. Here, we employ *in vitro* assays to investigate the impact of microtubule lattice defects on the severing characteristics of spastin. Toward this end, we prepared GMPCPP-stabilized microtubules with varying defect densities. Thereby, microtubule defects were introduced either through specific polymerization conditions or by end-to-end annealing of microtubules. We found that (i) the presence of defects accelerated the onset of the severing process and (ii) severing was twice as frequent in microtubule segments with defect sites as compared to random lattice segments. However, there was no evidence of preferential binding of spastin to defect sites. We therefore propose a severing mechanism in which defects do not actively promote microtubule severing but rather passively contribute to microtubule lattice instability, facilitating the process as fewer tubulin subunits are required to be removed for microtubule severing.

## Introduction

Microtubule regulators, including severing enzymes such as spastin, katanin and fidgetin organize the microtubule cytoskeleton inside cells by controlling their length and growth dynamics. Severing enzymes use the energy of ATP hydrolysis to extract tubulin subunits from microtubules and hence, generate internal breaks in the microtubule lattice which lead to severing events ^1, 2^. Microtubule severing by spastin and katanin plays pivotal roles in different cellular processes such as neurodevelopment ^3, 4^, mitosis and meiosis ^5, 6^. Furthermore, spastin contributes to endosomal fission and trafficking, shaping of the endoplasmatic reticulum ^7–9^ as well as controlling lipid droplet dispersion ^10, 11^ whereas katanin is heavily involved in ciliogenesis ^12^, cell migration ^13^ and in coordinating the remodeling, orientation and amplification of microtubule arrays in plants ^14, 15^. The diverse roles of microtubule severing, which involve both assembly and disassembly of the cytoskeleton, hint at severing enzymes having broader tasks than just severing microtubules.

Several studies have gained insights into the interaction and coordination of severing enzymes: Using catalytic inactive mutants, it was shown for spastin and katanin that the assembly of the enzymes into ring-shaped hexamers is needed for severing ^16, 17^. Unlike most AAA ATPases, katanin and spastin are monomeric when bound to ADP, and form hexamers in the presence of ATP ^16, 18, 19^. The assembly process for hexamer formation is not very well understood, however for spastin it has been postulated using bulk biochemical assays that the dimerization step, promoted by the monomer interaction with microtubules, is crucial in the formation of the active hexameric complex ^20^. It is thought that the monomers bind to the microtubules via electrostatic interactions, diffuse along the filament and oligomerize upon encounter with other monomers. Moreover, the C-terminal tails of tubulin were found to bind to the central pore of spastin and katanin hexamers ^17, 21, 22^. In this configuration the pull-out of a tubulin subunit from the lattice is supposed to occur due to conformational changes triggered by ATP hydrolysis. Importantly, fluorescence microscopy studies by Vemu et al. revealed that spastin and katanin can induce nanoscale damage by extracting a few tubulin heterodimers before actual severing occurs ^23^. This idea was supported by electron microscopy images and tomograms that visualized severing-mediated defects in which tubulin subunits were missing from the microtubule lattice *in vitro* and *in vivo* ^23, 24^. Interestingly, these damaged sites catalyzed the insertion of GTP-tubulin dimers from solution into the microtubule lattice, leading either to repaired filaments with GTP-islands or to rescued, stabilized microtubule fragments. Thus, severing enzymes may serve as quality control system for microtubules by removing dysfunctional tubulin and rejuvenating the microtubule lattice by incorporating fresh GTP-tubulin.

Supporting the latter hypothesis, it was suggested that the severing enzyme katanin preferentially severs at the irregularities of the microtubule lattice: (i) Computer modeling of microtubule severing in vitro could only replicate experimental results when lattice defects ^25^ (such as changes in protofilament number ^26^) were taken into account, and (ii) katanin was observed to localize and sever with increased frequency at the annealing sites between GMPCPP and GDP-taxol microtubules ^27^. Other studies have demonstrated that katanin has a strong affinity for microtubule crossovers and bundles: (i) In plants, severing by katanin at microtubule crossing sites resulted in alignment and reorientation of the microtubule cytoskeleton ^14, 15^, and (ii) in *Caenorhabditis elegans* oocytes, the pruning effect of katanin was confirmed and proposed to contribute to the maintenance of the (anti)parallel spindle architecture ^28^. In addition, katanin was found to be highly substrate-selective with respect to tubulin posttranslational modifications (PTMs) and microtubule-associated proteins (MAPs), preferentially severing older, more post-translationally modified microtubule parts ^29–31^. Thus, selective severing may also be a general mechanism for microtubule alignment and rearrangement.

So far, little is known about the mechanistic details of the severing process driven by spastin. While polyglutamylation, a type of tubulin PTM, has been shown to influence spastin activity ^32^, it is not clear whether spastin has the ability to recognize lattice defects or microtubule intersections. In order to gain deeper insight into the substrate dependence of spastin, we here studied how microtubule defects influence the severing process by spastin *in vitro*. Specifically, we induced defects in the microtubule lattice by either varying the polymerization conditions of microtubules or by end-to-end annealing of microtubules. Subsequently, microtubule severing by spastin was investigated with respect to defect densities and annealing sites. We found that the presence of defects significantly accelerated the onset of severing. Interestingly, our results do not indicate preferential binding of spastin to the defect sites leading us to propose a model where defects do not actively promote severing but rather passively contribute to microtubule lattice instability.

## Results

### Slow polymerization kinetics results in fewer microtubule-lattice defects

It has been shown that the polymerization conditions of microtubules influence the density of defect sites along the microtubule lattice, and hence, their mechanical stability ^33^. Therefore, we first investigated whether we could generate two distinct microtubule populations with different defect densities by varying the polymerization parameters. We tested two protocols to polymerize microtubules with the slowly hydrolyzing GTP-analog GMPCPP (**Figure 1A, Step I**, Methods): (i) For low defect density, microtubules were polymerized for 5 hours at low temperature (28 °C) at a moderate tubulin concentration of 2.5 µM (including 25 % rhodamine-labeled tubulin). We expected that the resulting slow microtubule assembly would lead to a low defect density. (ii) For high defect density, microtubules were polymerized for 30 min at high temperature (37 °C) at an elevated tubulin concentration of 20 µM (including 25 % Atto647-labeled tubulin). These conditions promote rapid microtubule assembly, expected to result in a high defect density.

**Figure 1.**
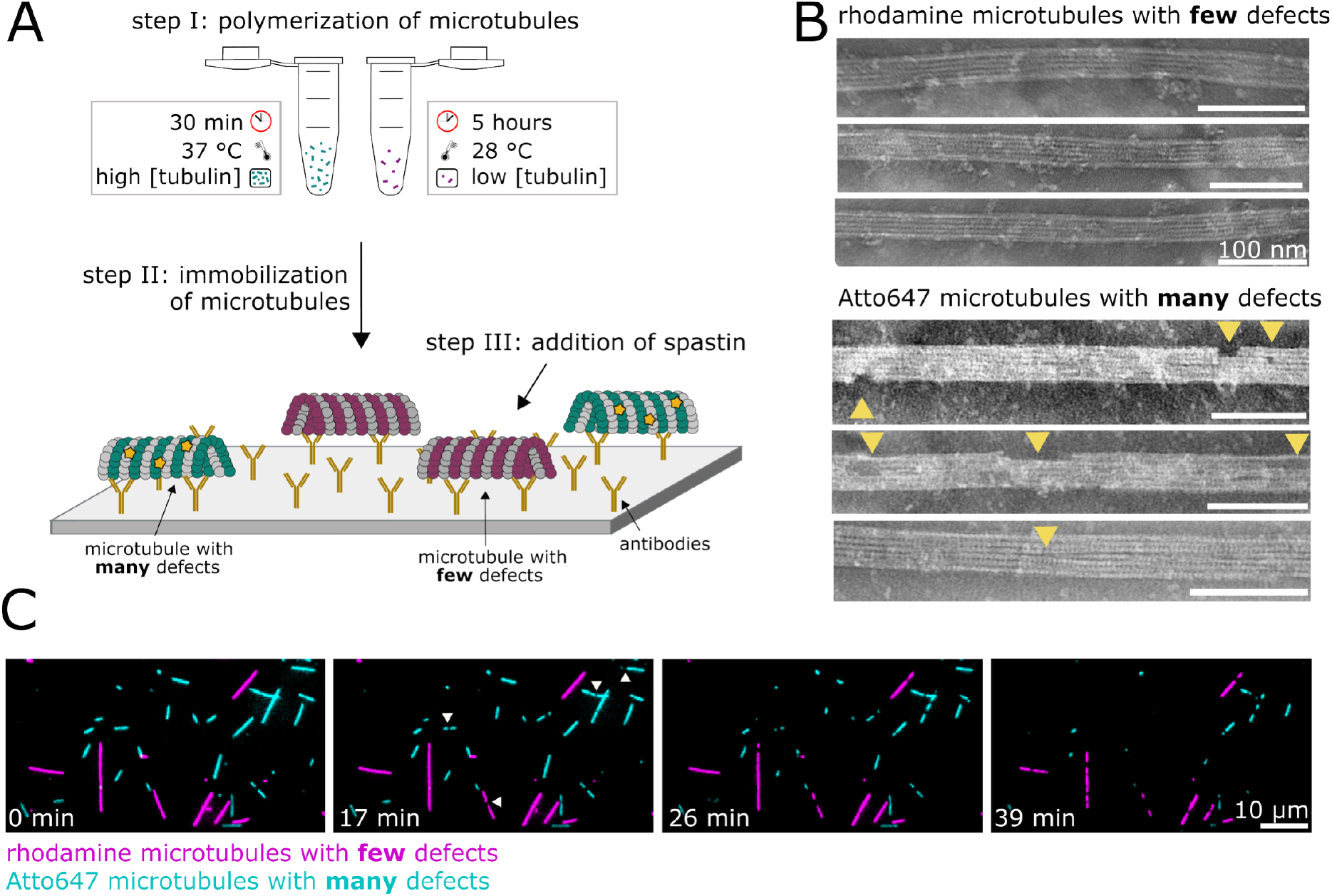
Severing of microtubules with few and many defects by spastin. **(A)** Schematic representation of the steps followed in a microtubule severing assay by spastin. Step I: Polymerization of microtubules according to two different protocols. Step II: Immobilization of microtubules with few and many defects onto a glass surface coated with anti-tubulin antibodies. Step III: Addition of the severing enzyme spastin. Green: Atto647-labeled tubulin; Magenta: Rhodamine-labeled tubulin. **(B)** Representative TEM images of microtubules with few defects (slowly-polymerized, at 28 °C for 5 hours with 2.5 µM rhodamine-labeled tubulin) and microtubules with many defects (rapidly-polymerized, at 37 °C for 30 min with 20 µM Atto647-labeled tubulin). Yellow arrows point at defect sites. (**C**) Fluorescence micrographs of Atto647 microtubules with many defects being severed faster than rhodamine microtubules with few defects (during a period of 39 min in the presence of 14 nM spastin). White arrowheads indicate the first severing sites of single microtubules.

In order to visualize the lattice structure of the microtubules resulting from the two polymerization protocols we applied transmission electron microscopy (TEM) (**Figure 1B**). We found that the lattices of slowly polymerized microtubules indeed hardly contained any visible defects. Moreover, no missing or broken pieces of microtubules were present. In contrast, the rapidly polymerized microtubules exhibited many different types of defects: missing or broken microtubule pieces in the lattice, changes in protofilament number and open structures due to disrupted lateral interactions. To ensure that the difference in defect density was due to the polymerization conditions and not because of differently labelled tubulin we also prepared slowly polymerized microtubules labeled with Atto647 and rapidly polymerized microtubules labeled with rhodamine (dyes exchanged) and obtained similar results (**Supplementary Figure 1**). The average defect densities were determined to be 1.46 ± 0.43 defects/μm (total analyzed microtubule length of 129 µm, N = 3 independent experiments) for rapidly polymerized microtubules versus 0.27 ± 0.09 defects/μm (total analyzed microtubule length of 126 µm, N = 4 independent experiments) for slowly polymerized microtubules. Thus, microtubules with two different defect densities (referred to below as microtubules with few or many defects) were successfully grown *in vitro* using different polymerization conditions.

### Severing of microtubules with many defects starts earlier than severing of microtubules with few defects

To investigate whether the density of defect sites in the microtubule lattice influences the onset and/or rate of severing by spastin, we immobilized microtubules with few defects and microtubules with many defects simultaneously on a glass coverslip in one flow channel using anti-tubulin antibodies (**Figure 1A, Steps II and III**). After addition of 14 nM spastin along with 1 mM ATP we followed the severing process for both types of microtubules using dual-color TIRF microscopy (**Figure 1C**). We observed an initial lag phase, during which the microtubules stayed intact, before severing events were detected at distinct positions of the microtubules. The severing process proceeded until the resulting microtubule fragments were so small that they detached from the surface and could not be reliably followed under the TIRF microscope. Comparison of the two types of microtubules showed that the severing onset of microtubules with many defects occurred earlier than of microtubules with few defects: the average time of the first ten severing events was 5.1 min and 13.4 min, respectively. The percentage of total microtubule length remaining at 30 min after addition of spastin was 37 % for microtubules with many defects compared to 70 % for microtubules with few defects.

In order to quantitatively compare the severing process for both microtubule populations (assay conditions as described above for **Figures 1A and 1C**), we counted the number of severing events that were observed during small time intervals (30 - 60 s) and added them up to plot their cumulative distribution. We found that severing started (and finished) earlier for microtubules with many defects as compared to the microtubules with few defects, while the severing progress itself was similar and took about the same time from its onset to the complete severing for both microtubule populations (**Figure 2A**).

**Figure 2.**
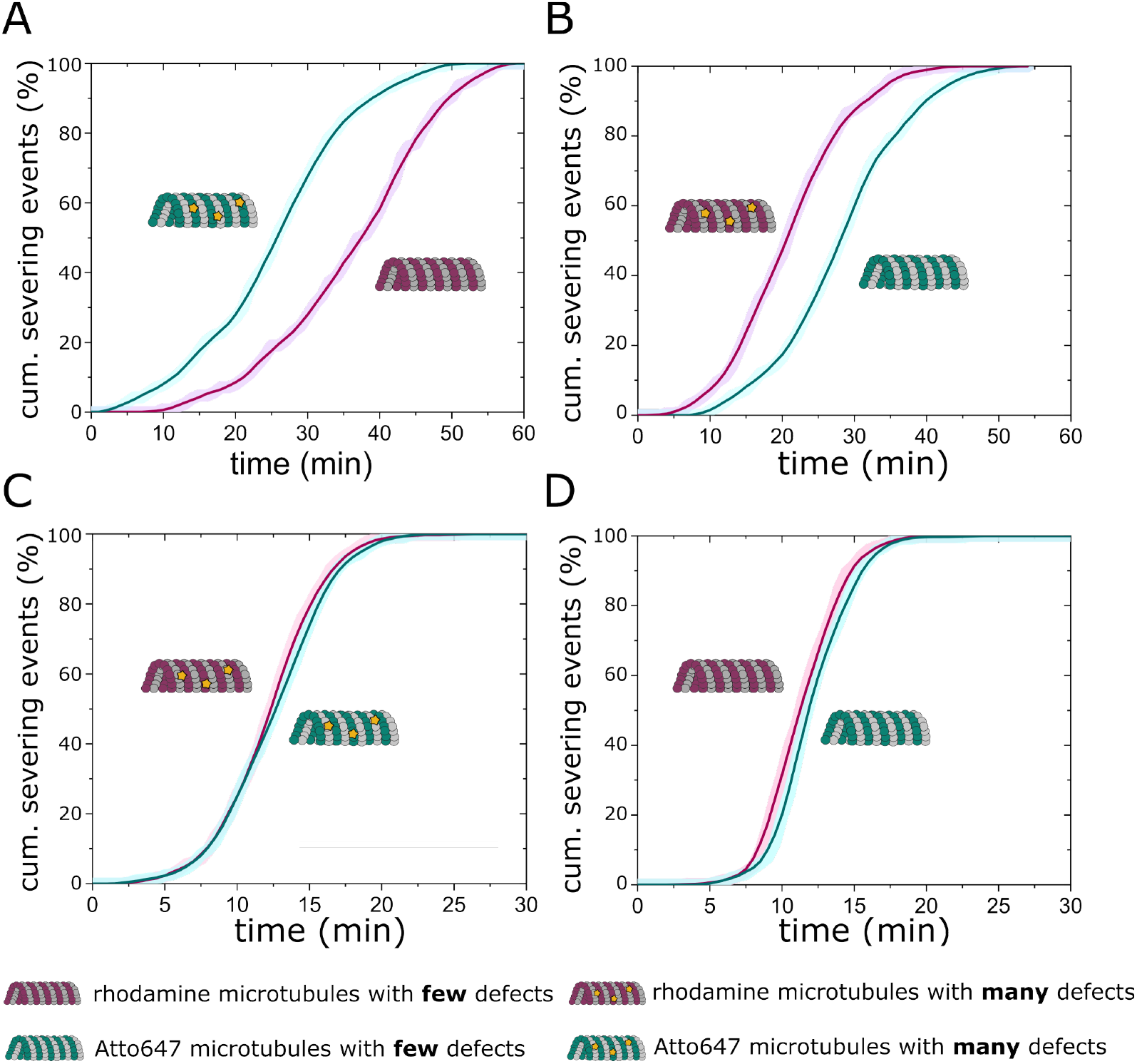
Severing time courses of microtubules with different fluorescent tags and defect densities. Cumulative severing events of two microtubule populations as a function of time at a spastin concentration of 14 nM (A, B) and 20 nM (C, D): **(A)** Rhodamine microtubules with few defects (N = 77; total length = 470 µm) and Atto647 microtubules with many defects (N = 135; total length = 483 µm) (corresponding assay in Figure 1C). **(B)** Rhodamine microtubules with many defects (N = 163; total length = 531 µm) and Atto647 microtubules with few defects (N = 95; total length = 573 µm). **(C)** Rhodamine (N = 388; total length = 1109 µm) and Atto647 microtubules (N = 297; total length = 1054 µm) with many defects, and **(D)** Rhodamine (N = 125; total length = 1029 µm) and Atto647 microtubules (N = 185; total length = 1101 µm) with few defects (corresponding assays in Supplementary Figure 2).

To exclude the possibility that the fluorescent labeling of the tubulin had an influence on the severing process, we performed a number of control measurements. First, we repeated the experiments described above by swapping the fluorophores on the microtubules with many and few defects. We again observed that the microtubules with few defects (now labeled with Atto647) could be observed longer in the field of view than the microtubules with many defects (now labeled with rhodamine, **Supplementary Figure 2A and Suppl. Movie 1**). The average time of the first ten severing events was 6.3 min and 11.0 min for microtubules with many and few defects, respectively. Furthermore, the percentage of total microtubule length remaining at 30 min after the addition of spastin was 19 % for microtubules with many defects compared to 55 % for microtubules with few defects. Likewise, the cumulative severing events over time (**Figure 2B**) showed again that severing started and finished earlier for the microtubules with many defects than for the microtubules with few defects, while the severing progress itself remained similar. As a second control, rhodamine- and Atto647-labeled microtubules, each either with few or many defects, were immobilized together in one flow channel. Upon addition of spastin (20 nM) in these assays, both types of microtubules were severed simultaneously, being present for the same amount of time (**Supplementary Figures 2B and 2C**), and no differences were found in the severing progress (**Figure 2C and D**). We therefore conclude, that the observed differences in severing between microtubule populations polymerized under different conditions can be attributed to the defect densities rather than the fluorescent tags on the tubulin.

### Microtubules are preferentially severed at annealing sites

To investigate whether severing preferentially occurs at defect sites, as compared to other parts of the microtubule, we annealed rhodamine- and Atto647-labeled GMPCPP microtubules with few defects over night at 30 °C (**Figure 3A, Steps I and II**). End-to-end annealing of microtubules has previously been reported to create microtubule lattice defects at annealing sites ^27, 33^. Importantly, the dual-color labeling of the microtubules allowed us to localize these defects. To test whether the annealed microtubules were reasonably stable and did not break apart at the annealing site in the absence of spastin, we performed gliding motility assays on substrate-immobilized kinesin-1 motors (**Supplementary Figure 3 and Suppl. Movie 2**). We found that the annealed microtubules were gliding smoothly without breaking, indicative of their overall mechanical stability in spite of annealing. Next, we immobilized annealed microtubules on the surface using anti-tubulin antibodies, added 25 nM spastin along with 1 mM ATP, and followed the severing process via two-color TIRF microscopy (**Figure 3A, Steps III and IV**). Severing events could be observed along the lattices of the annealed microtubules as well as at the annealing sites (**Figure 3B**). To compare the number and timing of severing events between annealing and random lattice sites, we divided the microtubules into 3 µm regions positioned either around the annealing site or entirely within the rhodamine-labeled or Atto647-labeled microtubule parts (**Figure 3C**). The 3 µm size of these regions was chosen because defects in the microtubule lattice likely not only influence the stability at the annealing sites themselves but also in the surrounding regions ^33^. The severing events were quantified by counting them for each region category for all microtubules in a field of view. For the representative microtubule in **Figure 3C**, two severing events were observed within the annealed region (region 2, white arrowheads) and one severing event in the non-annealed rhodamine region (region 1, yellow arrowhead). In a population of 68 microtubules we found a two-fold increase in the severing events per μm length of microtubule, during the observation period of 5 minutes, for regions with annealing sites compared to non-annealed regions with either rhodamine or Atto647 label (**Figure 3D**). No significant differences were observed in the severing events per μm length of microtubule between the rhodamine- and Atto647-only regions. Furthermore, we compared the timepoints of the first severing event within each of the 3 µm regions (**Figure 3E**). We found that the severing events in regions around annealing sites occurred at earlier timepoints than in rhodamine- or Atto647-only microtubule regions. These results are consistent with our previous observations, where the onset of microtubule severing occurred earlier for microtubules containing many defects as compared to microtubules containing few defects, independent of the fluorescent tag.

**Figure 3.**
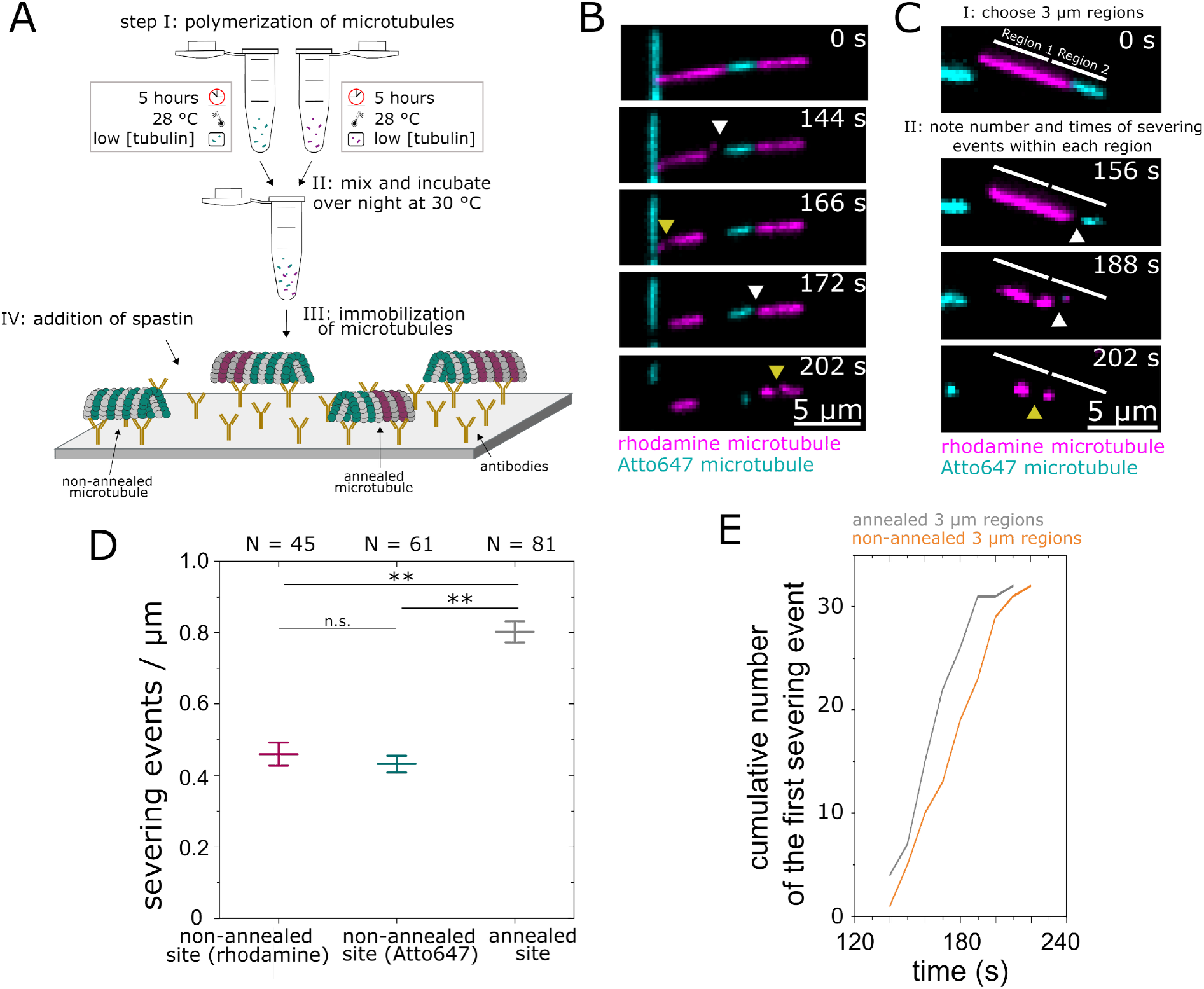
Severing of end-to-end annealed microtubules by spastin. **(A)** Schematic representation of the steps involved in the severing assay. Step I: Polymerization of differently labeled microtubules with few defects. Step II: Annealing of microtubules over-night. Step III: Immobilization of annealed microtubules onto a glass surface coated with anti-tubulin antibodies. Step IV: Addition of the severing enzyme spastin (25 nM). Green: Atto647-labeled tubulin; Magenta: Rhodamine-labeled tubulin. **(B)** Time course of a severing assay, in which a representative annealed microtubule is being severed at its annealing sites (white arrowheads) and at different parts of the microtubule lattice (yellow arrowheads). **(C)** Analysis steps followed to quantify the severing by spastin at annealing sites compared to random microtubule lattice sites. Step I: Microtubules were divided into 3 μm regions without (region 1: labeled with rhodamine) and with an annealing site (region 2). Step II: The number and timepoints of the severing events were determined for each region. **(D)** Severing events per μm (mean ± StDev) quantified for rhodamine-only, Atto647-only or annealed 3 μm microtubule regions. N = number of microtubules, time period of observation = 5 min. *P*-values were obtained by the two-sample T-test: **, *p* < 0.01. **(E)** Cumulative number of the first severing event within each 3 µm region as a function of time. An equal number of annealed and non-annealed regions were compared with each other.

### Tubulin loss and spastin signal evolve equally for microtubules with few and many defects

To test if tubulin loss and spastin signal develop over time in dependence of the defect density, we simultaneously imaged Atto647-microtubules (with many defects) and rhodamine-microtubules (with few defects) in the absence (control) and presence of 7 nM GFP-spastin (**Figure 4A**). In the absence of spastin the total microtubule length (sum of the lengths of all rhodamine-labeled microtubules (n=8) at a given time) only decreased slightly (less than 0.3% per minute) due to slow inherent end-depolymerization (**Figure 4B, upper panel**). In the presence of spastin, the total microtubule length (sum of the lengths of all microtubule parts remaining at a given time) first decreased slowly (similar to the case without spastin) but accelerated significantly in a non-linear manner after the onset of severing. The accelerated decrease in total microtubule length started earlier for microtubules with many defects compared to microtubules with few defects, corresponding to an early severing onset at 24.0 min (average time of first severing event per microtubule, N = 9 microtubules, cyan dotted line) and a late severing onset at 27.5 min (N = 9 microtubules, magenta dotted line), respectively. The length-averaged tubulin signal (fluorescence intensity per µm of microtubule) only slightly decreased in the absence of spastin (control) likely due to photobleaching (**Figure 4B, middle panel; Supplementary Figure 4**). In the presence of spastin, the length-averaged tubulin signal decreased significantly faster. Strikingly, this decrease started already before the onset of severing (as defined above, see dotted lines) indicating that tubulin removal already happens before actual severing is observed. We did not observe a significant difference in the decrease of length-averaged tubulin signal for microtubules with few and many defects. At the same time, the length-averaged spastin signal increased (**Figure 4B, lower panel; Supplementary Figure 5**). The persistent increase in the spastin signal over time likely reflects the oligomerization process of spastin, which is expected to lead to extended dwell times on the microtubule due to cooperative binding and/or lower off-rates after oligomerization. We did not observe a significant difference in time course and intensity of the spastin signals for microtubules with few and many defects. These results show that at the level of whole microtubules, there is no significantly increased or accelerated binding of spastin to microtubules with many defects compared to microtubules with few defects.

**Figure 4.**
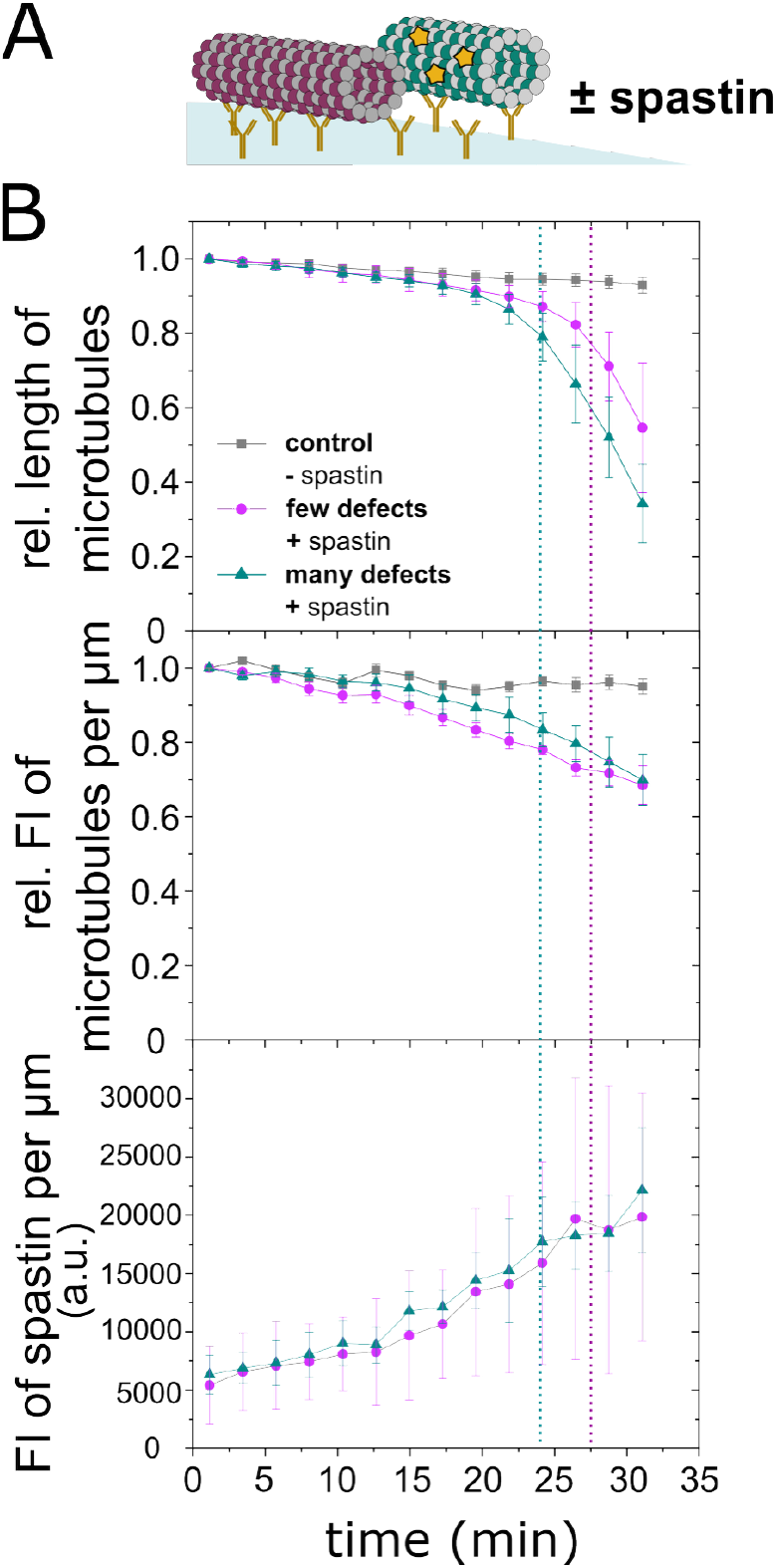
Comparison of the fluorescence intensity of GFP-spastin and tubulin for microtubules with different defect densities in dependence of time. **(A)** Schematic of the experimental assay, in which rhodamine-labeled microtubules with few defects were immobilized next to Atto647-labeled microtubules with many defects in the absence and presence of 7 nM GFP-labeled spastin. **(B)** Total microtubule length (relative to the value at t=0, ± StDev), fluorescence intensities (FI) of microtubules (relative to the value at t=0, ± StDev; Mann Whitney U-test at 8, 10.3, 19.6 and 21.9 min p > 0.01 but at all other time points *p* > 0.05 (n=7), see also Supplementary Fig. 4 for more data) and spastin (± StDev; Mann Whitney U-test at all time points *p* ≥ 0.086 (n=7)) per µm length of the remaining microtubule as functions of time. Dotted lines indicate the average time of the first severing event per microtubule (magenta - for microtubules with few defects; cyan - for microtubules with many defects).

## Discussion

In order to study the influence of lattice defects on microtubule severing by spastin, we employed two different tubulin polymerization conditions allowing for fast and slow microtubule growth kinetics, resulting in the generation of microtubules with many or few lattice defects, respectively. Using controlled *in vitro* severing assays, we present experimental evidence that a higher number of defects in the microtubule lattice results in an earlier onset of the severing process by spastin. Furthermore, we generated localized defect sites by end-to-end annealing of slowly-polymerized microtubules. On these microtubules, severing occurred twice as often in the annealed regions after 5 min compared to other regions of the microtubule lattice with presumably less defect sites. The latter result is quantitatively comparable to findings made with katanin, which has been reported to sever microtubules with an about 50 % higher frequency at annealing sites between GMP-CPP and taxol-stabilized GDP microtubules, compared to regions away from the annealing sites ^27^. Because in those experiments, the two microtubule types are polymerized in the presence of different nucleotides, they are likely not only conformationally different with different protofilament numbers but may also contain different defect densities and hence both these factors can influence the severing frequency. To rule out such influences, we consistently employed GMP-CPP stabilized microtubules. In those earlier experiments, the katanin molecules were observed to quickly localize to microtubule annealing sites and remained there until severing had taken place^27^. In contrast, in our experiments spastin bound rather uniformly along the microtubule lattice, and even at low spastin concentrations (**Supplementary Figure 5**) we did not observe an obvious accumulation of GFP-spastin at particular sites on the microtubule lattice.

What causes microtubules (or regions of them) with many defects to be severed earlier than microtubules (or regions of them) with few defects? Let us assume the following established mechanism of microtubule severing by spastin: Spastin hexamers (which form on the microtubule lattice) remove tubulin dimers one after another from the microtubule lattice ^34^. Once a certain number of tubulins are removed in close vicinity (i.e. once a given threshold density of missing tubulins in the microtubule lattice is reached locally) the microtubule is severed at this location. In the case of lattice defects, two main mechanisms may be at play. (i) **Defect recognition model:** Spastin monomers and/or hexamers directly recognize defect sites in the microtubule lattice (i.e. they bind/interact there with higher affinity compared to the rest of the lattice). This leads to a local increase in spastin concentrations, resulting in higher tubulin removal rates at the defect sites. (ii) **Lattice instability model:** Spastin activity on the microtubule lattice is homogeneous along the length, i.e. tubulin removal occurs at random sites. However, at lattice defect sites where tubulins are already missing and the lattice stability is compromised, fewer tubulins need to be removed for severing to occur. For both models, severing would occur earlier on microtubules with many defects compared to microtubules with few defects because the threshold density of locally missing tubulins is reached faster. Additional synergistic effects are expected for the defect recognition model, because at the defect sites, with missing tubulin subunits, fewer tubulins have to be removed by spastin to reach the critical density required for severing. Moreover, for both models severing would be expected to occur predominantly in the vicinity of (pre)existing defect sites (either present already at the beginning or generated by spastin itself). While we cannot make a statement about this correlation for our homogeneously grown microtubules with few and many defects (as we do not have the knowledge about the exact positions of the defect sites), we do know from our experiments with annealed microtubules and from ref. ^27^ that severing indeed occurs preferentially near defect sites. Beyond these commonalities, the defect recognition model predicts a higher and rapidly increasing spastin signal, accompanied by a faster decline in the tubulin signal prior to severing, in microtubules with many defects compared to few defects. However, this was not observed in our experiments (**Figure 4B, middle and lower panels**). Instead, our experimental data indicate that the average tubulin and spastin signals are independent of defect density, supporting the lattice instability model.

The results of our study suggest that spastin molecules bind randomly to microtubules and remove tubulin subunits without preferential binding or increased affinity for defect sites. Spastin could either create new defects or enlarge existing ones with similar probability. Moreover, the need to remove less tubulin leads to an earlier onset of severing. The continuous severing progress with similar severing rates for both microtubule types (**Figure 2**), even after an earlier onset for microtubules with many defects, indicate that microtubules with few defects behave similarly to those with many, as spastin generates more “defects” in both. The preference of spastin for specific sites or local accumulations observed *in vivo* ^35^ could be due to factors such as localized microtubule posttranslational modifications and MAP decoration ^32, 36–40^. Some of these factors could “mark” microtubule lattice defects leading to preferential severing at these sites – a mechanism also proposed by Kuo et al. ^34^ as an explanation for the strong affinity of katanin for microtubule crossovers and bundles *in vivo* ^14, 15, 29, 41^. Alternatively, the presence of posttranslational modifications and MAPs could specifically generate defects at these sites, enabling localized microtubule polymerization and the creation of new minus ends ^23, 35, 42, 43^.

## Materials and Methods

### Preparation of flow channels

Experiments were performed in 3 mm wide flow channels composed of a dichlorodimethylsilane (DDS)-coated coverslip^44^ (22×22 mm^2^, Menzel-Gläser) at the bottom, a polyethylene glycol (PEG)-coated coverslip^45^ (18×18 mm^2^, Menzel-Gläser) on top and two stripes of parafilm as spacers in between by heating the flow sample to 80 °C on a hot plate. For gliding motility assays, glass coverslips (22×22 mm^2^, Menzel-Gläser), which were cleaned according to the following procedure, were used at the bottom of the flow channel: Sonication in Mucasol/water (1 : 20; v/v) for 15 min was followed by rinsing in deionized water for 2 min. Further, coverslips were sonicated in ethanol/water (1:1; v/v) for 10 min, rinsed in deionized water for 2 min, rinsed in MilliQ-water for 2 min and finally dried using a nitrogen airflow.

### Spastin purification

A human spastin plasmid short isoform (Δ227)^19^ was cloned into Optimized Classic Cloning vectors developed by Lemaitre et al. (2019)^46^ to produce two different constructs: (1) containing a recombinant N-terminal 6x His-tag and (2) containing a N-terminal 6x His-tag followed by a GFP-tag for fluorescently labeling. The vectors were expressed in E.coli cells (BL21, pRARE competent cells). After transformation, cells were pelleted and resuspended in Lysis buffer (50 mM sodium phosphate buffer, 300 mM KCl, 5 % Glycerol, 1 mM MgCl_2_, 10 mM β-mercaptoethanol, 0.1 mM ATP, pH 7.4), with 30 mM Imidazole, and protease inhibitor cocktail (cOmplete, EDTA free, Roche). All following steps were performed at 4 °C. The cells were lysed by flowing the resuspended cell solution through an Avestin Emulsiflex three times. The cell lysate was then centrifuged for 45 min at 40,000 rpm using Type 45 Ti rotor in an Optima XPN-80 ultracentrifuge following the addition of Benzonase (1:10000, from Sigma Aldrich). The supernatant was filtered through a 0.45 µm membrane and subsequently incubated with Ni-NTA agarose beads (160019892, Qiagen) for 2 h. The beads were washed several times in a 10 ml gravity flow column (Econo-Pacchromatography columns, Biorad) with equilibration buffer (Lysis buffer with 30 mM Imidazole) and eluted with equilibration buffer containing 200 mM imidazole. The 6x His-tag was cleaved over night with PreScission 3C-protease. The uncleaved fraction as well as the His-tagged protease was subsequently removed by incubating with Ni-NTA resin, and the flow-through containing cleaved spastin fraction was concentrated using 10 K MWCo spin filter, snap-frozen in liquid nitrogen and stored at −80 °C.

### Kinesin-1 purification

A wild type kin-1 construct consisting of full length Drosophila melanogaster kin-1 heavy chain with a C-terminal 6x His-tag was expressed and purified as described earlier^47^.

### Tubulin purification

Porcine tubulin was purified from porcine brain using an established protocols^48^. Tubulin was fluorescently labeled with either rhodamine or Atto647N in BRB80 with 5 mM MgCl2, 1 mM GTP, 5 % Dimethyl Sulfoxide (DMSO) at 37°C for 30 min. The ratio of labeled to unlabeled tubulin was 1:3.

### GMPCPP-stabilized microtubules

Two slightly different protocols were used to grow microtubules with few and many defects, respectively. Microtubules with many defects were polymerized by mixing 20 µM rhodamine- or Atto647N-labeled tubulin with 2 mM GMPCPP and 0.1 mM MgCl2 in Brinkley Reassembly Buffer 80 mM (BRB80, adjusted to pH = 6.9 with KOH and composed of 80 mM 1,4-piperazinediethanesulfonic acid (PIPES), 1 mM EGTA, and 1 mM MgCl2) in a final volume of 10 µl followed by an incubation step of 30 min at 37 °C in a dry block heater. Finally, 190 µl BRB80 were added to the microtubule solution that was then centrifuged in an airfuge for 5 min at 150,000 x g and resuspended in 150 µl of BRB80. Microtubules with few defects were polymerized with 2.5 µM rhodamine- or Atto647-labeled tubulin, 1.25 mM GMPCPP, and 1.25 mM MgCl2 in BRB80 in a final volume of 80 µl for 5 hours at 28 °C in a dry block heater. Finally, 120 µl BRB80 were added to the microtubule solution that was then centrifuged in a tabletop centrifuge at 17,000 x g during 15 min and resuspended in 150 µl of BRB80.

### Preparation of end-to-end annealed microtubules

In order to anneal differently labeled GMPCPP microtubules, rhodamine and Atto647-labeled microtubules were polymerized separately by following the protocol for microtubules with few defects. Then, equal volumes of rhodamine and Atto647 microtubules were mixed and incubated over night at 30 °C. Before starting the experiment, the microtubule mixture was pelleted in a tabletop centrifuge at 17,000 x g for 15 min and resuspended in 150 µl of BRB80.

### Microtubule severing assays

First, flow cells were flushed with 20 µl of the following sequence of solutions to immobilize microtubules on the surface: (i) 1500x dilution of TetraSpeck microspheres (0.1 µm diameter, lot#1786280 Thermo Fisher Scientific) to allow drift correction; incubation time 5 min, (ii) solution of monoclonal anti-β-tubulin I mouse antibodies; incubation time 5 min, (iii) twice BRB80 for washing, (iv) BRB80C buffer (0.45 mg/ml Casein in BRB80) for blocking the surface from unspecific protein adsorption; incubation time 5 min, (v) dilution of GMPCPP-stabilized microtubules in BRB80; incubation time 10 min, and (vi) antifade solution (110 µg/ml glucose oxidase and 22 µg/ml catalase in Hepes based buffer). Then, for severing the immobilized microtubules, spastin ([spastin] = 24 µM; [GFP-spastin] = 34.3 µM) was first diluted 10x - 40x in Hepes buffer (20 mM Hepes, 75 mM KCl, 0.5 mg/ml Casein, 0.1 % Tween-20, 1 mM ATP, 20 mM glucose, 10 mM DTT) and then further in antifade solution in Hepes buffer to achieve the final concentration indicated for each experiment. The spastin dilution was then perfused into the flow cell, which was already placed on the microscope, and imaging was started immediately.

### Gliding motility gliding assays

The flow cell was first perfused with 20 µl of BRB80C buffer to coat the cleaned coverslip with casein. The solution was left to adsorb for 5 min. Then, 20 µl of a kinesin-1 solution (2 µg/ml kin-1, 10 mM DTT, 1 mM ATP and 0.2 mg/ml Casein in BRB80) were added and incubated for another 5 min. Thereafter, 20 µl of motility solution (1 mM ATP, 20 mM Glucose, 10 mM DTT, 33 nM microtubules, 56 µg/ml glucose oxidase and 11 µg/ml catalase in BRB80) containing microtubules was applied. After 5 min, unbound microtubules were washed out with antifade solution in BRB80 (1 mM ATP, 20 mM Glucose, 10 mM DTT, 20 µg/ml glucose oxidase and 10 µg/ml catalase in BRB80) and the gliding microtubules were imaged.

### Optical Microscopy

All the experiments were performed at 28 °C using an objective heater unless indicated otherwise. An inverted fluorescence microscope (Axio Observer Z1) with a 63x oil immersion 1.46 NA objective (Zeiss) and an EMCCD camera (Andor Technology) controlled by the software Metamorph 7.7.7 was used. A built-in magnification (1.6x Optovar) was additionally implemented leading to a pixel size of 0.159 µm. Rhodamine or Atto647-labeled GMPCPP-stabilized microtubules were imaged via epifluorescence or TIRF microscopy, indicated for each experiment. For epifluorescence illumination a Sola SE2 (13212) Lumen 200 metal arc lamp with a GFP, TRITC or Cy5 filter was used. TIRF imaging was performed by using the Omicron Laserbox (Rodgau-Dudenhofen, with the 488 nm, 561 nm and 642 nm lasers), a QuadLine TIRF filter (AHF, 405/488/561/640x) and adjusting the TIRF angle before starting with an experiment. A fast filter wheel (Finger Lakes Instrumentation) was employed to image two colors nearly simultaneously (time difference = 123 ms) in some of the experiments, as indicated. An exposure time of 100 ms was employed. The time between frames was 2 s in experiments in which the severing of microtubule with few and many defects was investigated over time. The fluorescence signal of GFP-spastin on microtubules was acquired in stream mode (10 images / s), and about every 130 s an image of the microtubules was taken. To determine the change in microtubule fluorescence intensity over time (**Supplementary Figure 4**), imaging was performed using a Nikon Eclipse Ti2 microscope equipped with a perfect focus system and a 100 ×, 1.49 NA oil, apochromat TIRF objective, and 1.5 × optovar. Samples were illuminated with 550 nm (75 %) and 660 nm (25 %) of a CoolLED pE-4000 lamp. Images from different fluorescent channels were acquired with separate EMCCD cameras (iXon Life EMCCD for 550 nm and iXon Ultra EMCCD for 660 nm), each containing 1024 × 1024 pixel sensor and controlled with VisiView. The size of each pixel was 87 × 87 nm. Images were acquired in time-lapse mode with 100 ms exposure (1 frames every 5 seconds).

### Transmission electron microscopy

Carbon-coated 400 mesh copper transmission electron microscopy (TEM) grids (S160-4 Plano, Wetzlar) were treated with plasma etching for 15 s to make the surface of the grid hydrophilic and enable the adhesion of microtubules. Then, after centrifugation, the pellet of microtubules with few and many defects, respectively, was resuspended in 100 µl BRB80. 5 µl of the resuspended microtubule solution were carefully placed on the grid and incubated for 10 min. The excess sample solution was absorbed by a clean paper towel from the edges of the grid. The grids were allowed to dry completely for 20 min. Finally, the grids were stained: To a 0.75 % uranyl formate solution prepared in advance and stored at −20 °C, sodium hydroxide solution was added at a final concentration of 25 mM. After quickly washing the dried grids with 10 µl of ultrapure water, 10 µl of the staining solution were applied for 15 s. Then, the stain was carefully removed with a clean paper towel and the grids were allowed to dry over-night. The TEM samples were imaged using a FEI Morgagni 268D transmission electron microscope operated at 80 kV in combination with an Olympus MegaView III camera of the CMCB Electron Microscopy Facility located at the Center for Regenerative Therapies Dresden.

### Data processing and analysis

Severing events (microtubule breaks) were counted manually as a function of time, from the beginning of the timelapse movie immediately after the addition of spastin until no more microtubules were in the field of view (*i*.*e*., when no further severing events were possible). Severing events were plotted as cumulative events normalized from 0 to 100. To quantify the number of severing events per μm microtubule length in regions around an annealing site compared to a random microtubule lattice site (labeled with either rhodamine or Atto647), we divided each microtubule into 3 μm-sized regions as shown in **Figure 3C** for a representative microtubule. For each type of region (rhodamine only, Atto647 only or annealed), the number and timing of severing events was determined. ImageJ software was employed for image processing. Fluorescence intensity data was acquired by tracking of microtubules with Fluorescence Image Evaluation Software for Tracking and Analysis (FIESTA)^49^ (**Supplementary Figure 4**), which automates Gaussian fitting of fluorescence signals (or in **Supplementary Figure 5** by line scans along microtubules using ImageJ software). Tracked data was manually corrected to exclude erroneous tracks. Since microtubule intensity data was only collected ~ every 130s, the average of the two time points before and after each stream of GFP-spastin was plotted at the same time point as the corresponding spastin signal.

## Acknowledgements

We thank Foram Joshi and the Electron Microscopy Facilities at the Center for Molecular and Cellular Bioengineering (CMCB) of TUD Dresden University of Technology for expert support with electron microscopy, Günther Woehlke for the kind donation of the spastin plasmid, Corina Bräuer for technical assistance and all members of the Diez laboratory for scientific discussions. This work was funded by the Technische Universität Dresden.

