## Supplementary Information for "Microtubule lattice defects facilitate spastin-mediated severing"

* shared, first authors

---

### Supplementary movies

**Movie S1. Severing assay of microtubules with different defect densities.** Microtubules labeled with Atto647 (cyan color) were slowly polymerized and thus contained only few defects. Rhodamine labeled microtubules (magenta color) were grown rapidly resulting in many defects. Both microtubules population were immobilized on the surface and the severing progress by spastin was followed over time.  Cyan-colored microtubules with few defects could be observed longer in the field of view than magenta microtubules with many defects.

**Movie S2. Gliding motility assay with annealed microtubules.** Microtubules, fluorescently labeled with either Atto-647 or rhodamine and annealed overnight, were gliding across a surface coated with kinesin-1 molecules. During the observation period none of the annealed microtubules broke apart.

### Supplementary Figures

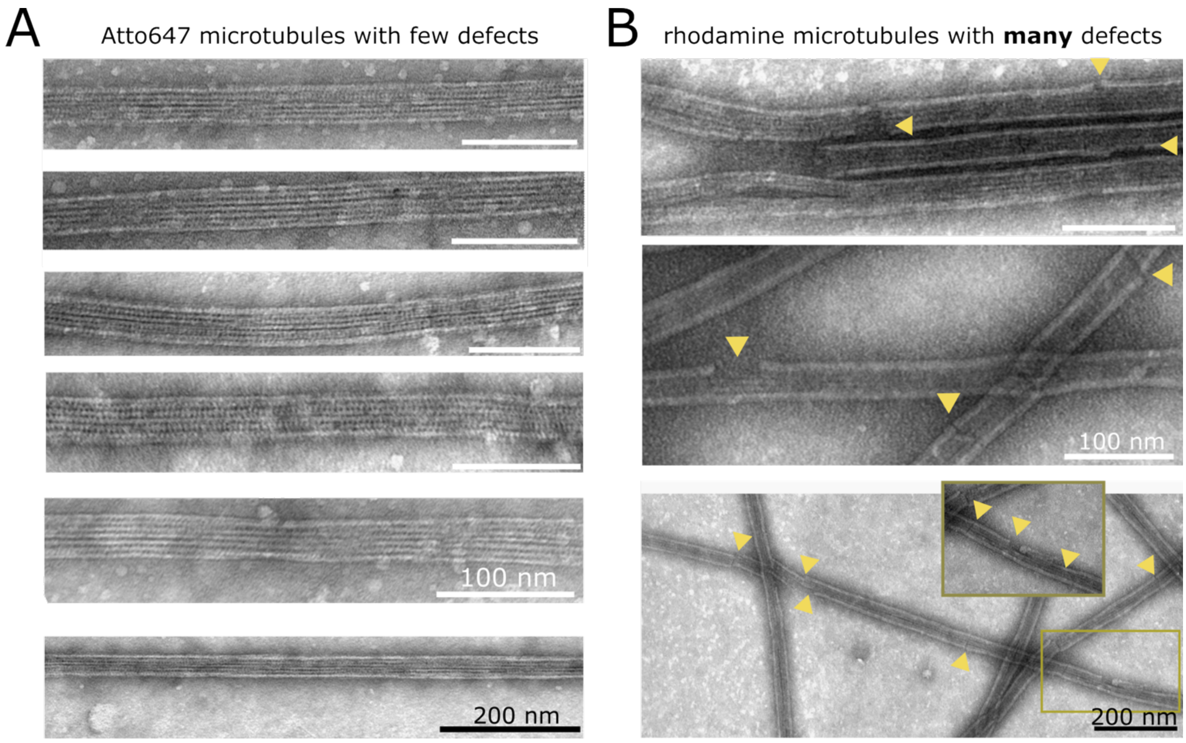


**Supplementary Figure 1. Representative TEM images of microtubules.** The microtubules in **(A)** were polymerized slowly at 28 °C for 5 hours with 2.5 µM Atto647-labeled tubulin and those displayed in **(B)** were polymerized rapidly at 37 °C for 30 min with 20 µM rhodamine-labeled tubulin. Compared to Figure 1B, the dyes for each polymerization condition are interchanged. Yellow arrows point to defect sites.

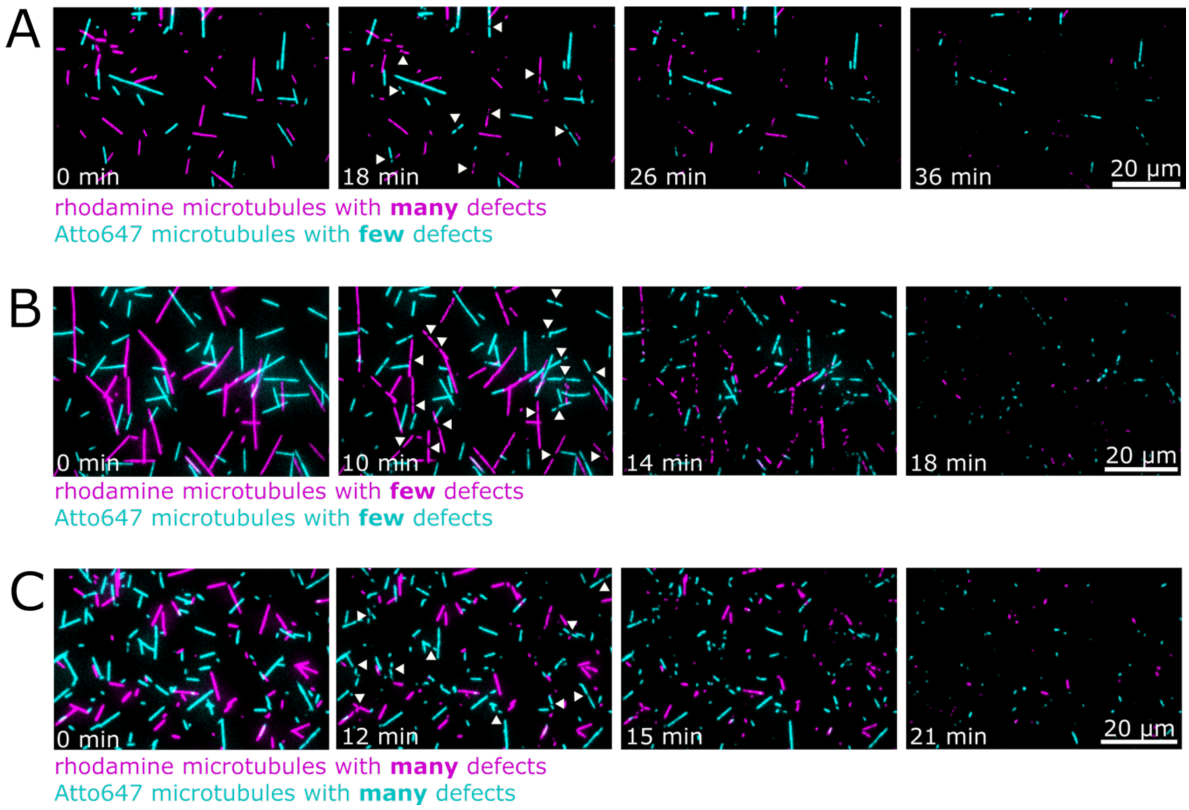


**Supplementary Figure 2. Severing assays of microtubules with different fluorescent tags and defect densities.** The time series of color overlays of fluorescence images in **(A)** shows that rhodamine-labeled microtubules with many defects were severed faster than Atto647-labeled microtubules with few defects (in the presence of 14 nM spastin). However, if microtubules with different fluorescent tags were polymerized under the same conditions, with few defects **(B)** or many defects **(C)**, they were severed in a similar time frame (in the presence of 20 nM spastin). White arrows indicate the first severing site of a single microtubule.

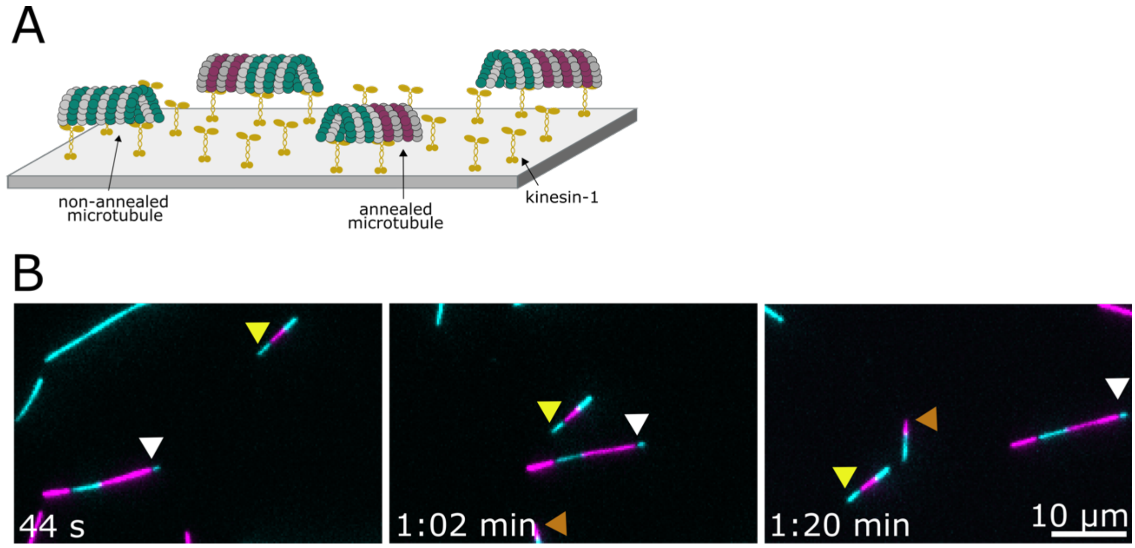


**Supplementary Figure 3. Gliding motility assay with annealed microtubules. (A)** Schematic representation of a microtubule gliding assay. Surface-attached kinesin-1 motor proteins propel, in the presence of ATP, microtubules with and without annealing sites across the surface. **(B)** Fluorescence images of a representative time sequence illustrating the movement of annealed microtubules. Arrowheads of different colors point to different microtubules.

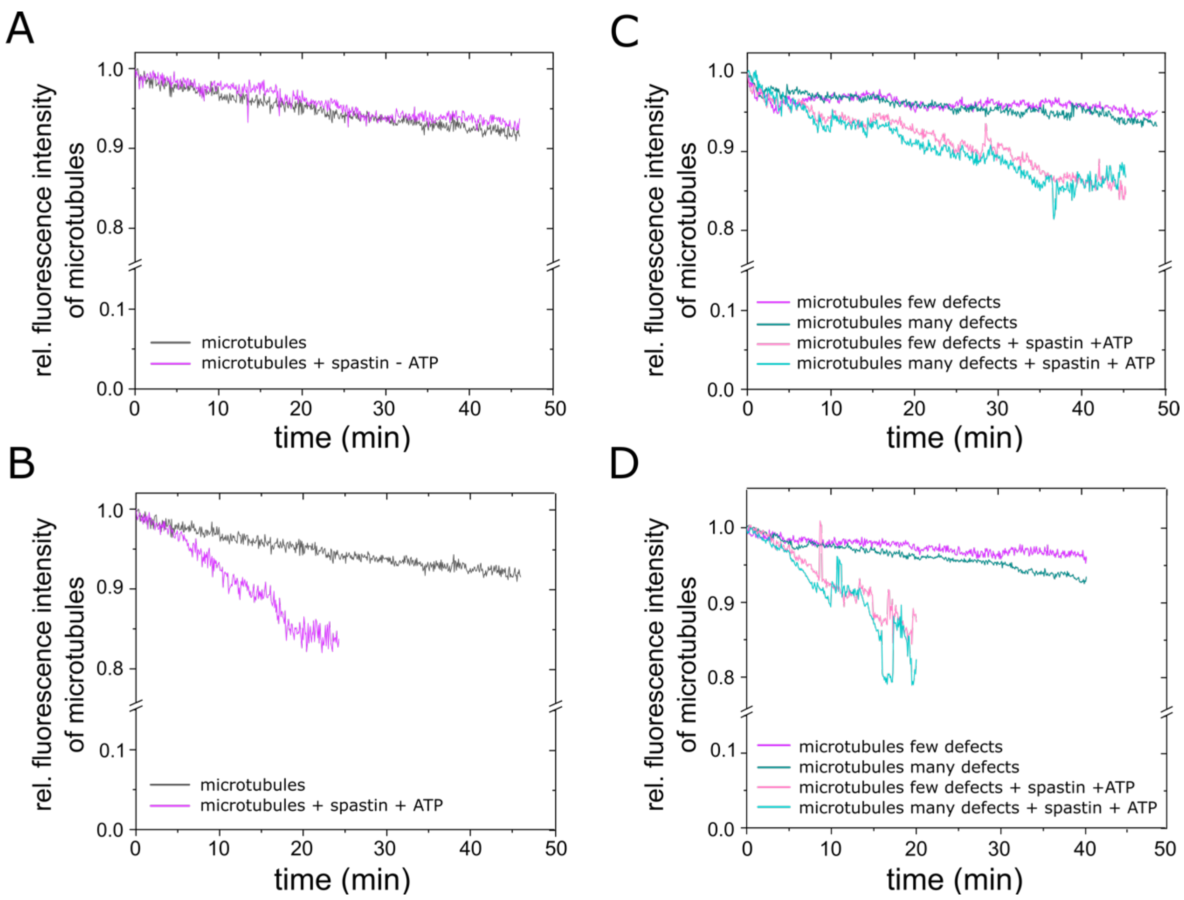


**Supplementary Figure 4. Change in microtubule fluorescence intensity over time in dependence of spastin concentration and activity. (A)** Relative average rhodamine fluorescence intensities per µm of microtubules without severing (grey: no spastin; magenta: 7 nM spastin but no ATP). **(B)** Relative average rhodamine fluorescence intensities per µm of microtubules without severing (grey: no spastin) and with severing (magenta: 7 nM spastin with ATP). **(C)** Relative average fluorescence intensities per µm of microtubules without severing (no spastin; magenta: rhodamine-labeled microtubules with few defects; green: Atto647-labeled microtubules with many defects) and with severing (3.5 nM spastin with ATP; pink: rhodamine-labeled microtubules with few defects; cyan: Atto647-labeled microtubules with many defects; Mann Whitney U-test at 18 time points (1 to 35 min): $p > 0.05$ (n=12)). **(D)** Same experiments with 7 nM spastin with ATP (Mann Whitney U-test for microtubules with spastin at 10 time points (1 to 19 min): $p > 0.05$ (n=14)). One image was acquired every 5 s with LED lamp (75 % / 25 % lamp intensity for 550 nm and 660 nm excitation, respectively; in D: 70 % / 60%). In (A) to (C) intensities were determined by tracking at least 12 microtubules with FIESTA and then averaging them. In (D) the microtubule lengths were determined by FIESTA and the intensities were measured with Fiji. For all values the background intensities (obtained from image areas next to the microtubules) were subtracted.

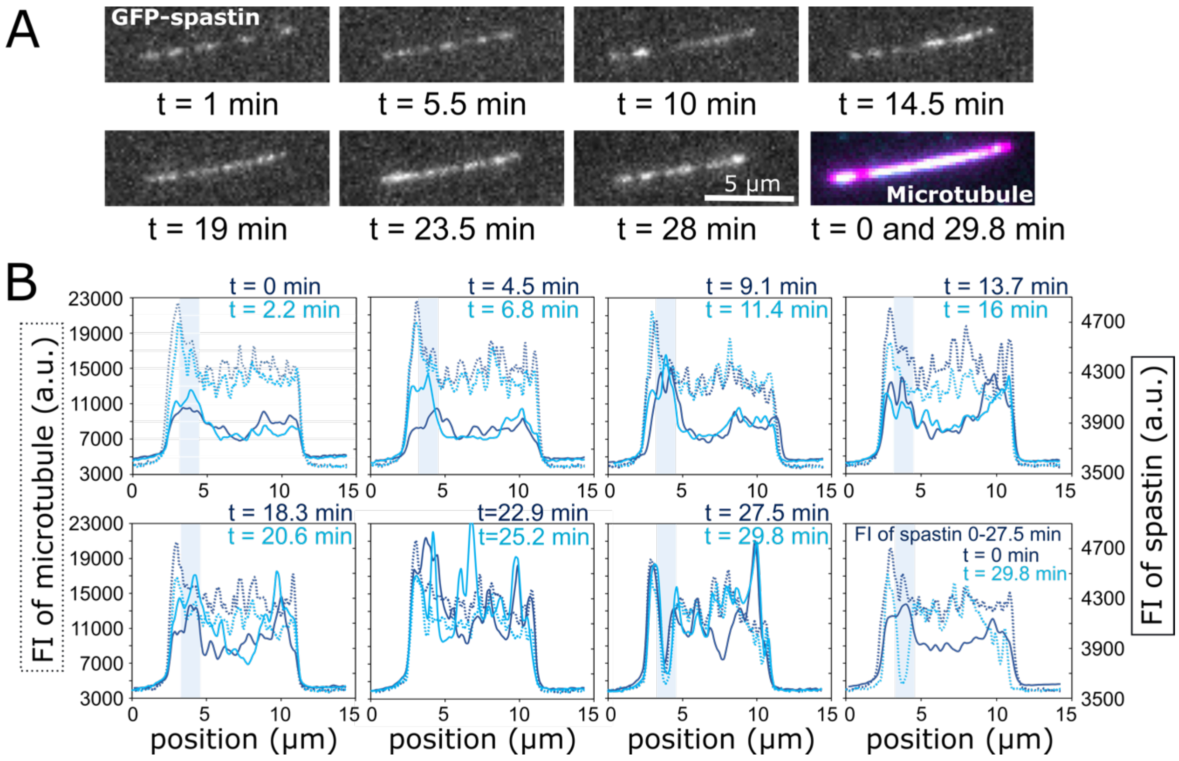


**Supplementary Figure 5. Fluorescence signal of GFP-spastin along a microtubule. (A)** Time series of fluorescence images of GFP-spastin on a microtubule at different time points with the same scaling of intensity. **(B)** Fluorescence intensity profiles along a rhodamine-labeled microtubule over the time-course of the experiment. Each diagram displays the average fluorescence intensity of spastin (solid lines) of two consecutive streams (each 1000 images) as well as the fluorescence intensity of the microtubule (dotted lines) before each of the two streams. In the last diagram the spastin intensity was averaged over stream 1-12 (0-27.5 min), the time period from the beginning of imaging until the first severing site became visible. Moreover, the microtubule intensity is shown before stream 1 and 14 (0 - 29.8 min). The severing position is indicated by a light blue bar.
